# The anti-apoptotic proteins Bcl-2 and Bcl-xL suppress Beclin1/Atg6-mediated lethal autophagy in polyploid cells

**DOI:** 10.1101/684266

**Authors:** Jing Zhang, Shenqiu Zhang, Fachi Wu, Qiong Shi, Thaddeus Allen, Fengming You, Dun Yang

## Abstract

Inhibition of Aurora-B kinase is a synthetic lethal therapy for tumors that overexpress the MYC oncoprotein. It is currently unclear whether co-occurring oncogenic alterations might influence this synthetic lethality by confering more or less potency in the killing of tumor cells. To identify such modifers, we utilized isogenic cell lines to test a variety of cancer genes that have been previously demonstrated to promote survival under conditions of cellular stress, contribute to chemoresistance and/or suppress MYC-primed apoposis. We found that Bcl-2 and Bcl-xL, two antiapoptotic members of the Bcl-2 family, can partially suppress the synthetic lethality, but not multinucleation, elicited by a pan-aurora kinase inhibitor, VX-680. We demonstrate that this suppression stems from the rescue of autophagic cell death, specifically of multinucleated cells, rather than a general inhibition of apoptosis. Bcl-2 inhibits VX-680-induced death of polyploid cells by interacting with the autophagy protein Beclin1/Atg6 and rescue requires localization of Bcl-2 to the endoplasmic reticulum. These findings expand on previous conclusions that autophagic death of polyploid cells is mediated by Atg6. Bcl-2 and Bcl-xL negatively modulate MYC-VX-680 synthetic lethality and it is the anti-autophagic activity of these two Bcl-2 family proteins, specifically in multinucleate cells, that contributes to resistance to Aurora kinase-targeting drugs.

## Introduction

There is a propensity for MYC oncogenes to be mutated and/or overexpressed and this contributes to the genesis of many types of human cancer [1] [2] [3]. The MYC family includes three members, MYC, MYCN, and L-MYC [3]. Collectively, this family of “super-transcription factors” has been found to be deregulated in over 50% of human cancers and is thought to regulate at least 15% of the entire human genome [2]. Hence many cellular processes, including ribosome biogenesis, protein translation, cell-cycle progression, and metabolism, are altered with deregulaiton of MYC transcription factors.

Many lines of scientific research support the supposition that targeting MYC can be an effective strategy in cancer therapy. However, direct targeting of MYC has proven technically difficult. Instead, a variety of synthetic lethal therapies have been developed to target tumor cells that overexpress MYC [4] [5] [6] [7]. Among them is a synthetic lethal therapy that allows the selective killing of cells that overexpress MYC by pharmacological inhibitors of the Aurora-B kinase (AURKB). Inhibitors have been tested clinically [8]. The lethality with AURKB inhibition is executed by an early phase of apoptosis and a delayed phase of autophagy that kills MYC-promoted polyploid cells in the absence of cytokinesis.

It is well known that certain genomic alterations serve to promote cell survival and are responsible for resistance to conventional chemotherapies, targeted therapies and immunotherapy [9] [10] [11]. Examples include mutation or deletion of the tumor suppressor p53, constitutive activation of RAS, AKT or PI3K and overexpression of Bcl-2, among other alterations. These are often found alongside deregulated expression of MYC in human malignancy. We hypothesized that these same chemotherapy resistance-promoting genes and pathways might also modify the synthetic lethal interaction between MYC and Aurora kinase inhibitors, thereby limiting clinical efficacy. We directly tested this hypothesis and by assaying the influence of various cancer genes on the MYC-VX-680 synthetic lethal interaction.

We modeled the impact of these oncogenic alterations using a panel of isogenic retinal pigment epithelial cells that constitutively overexpress MYC (RPE-MYC). In RPE-MYC cells, AURKB inhibition, or other manipulations that disable the chromosomal passenger protein complex, mediate synthetic lethity [5]. Disabling AURKB with VX-680 elicits a bimodal cell death characterized as early apoptosis followed by delayed lethal autophagy. Our system enabled us to distinguish if a defined oncogenic element could impact either one or both phases of cell death. We discovered that the proapoptotic members of the Bcl-2 family, Bcl-2 and Bcl-xL, suppressed autophagic, not apoptotic, cell death elicited by the synthetic lethal interaction. Suppression could be attributed to the interaction between either of the two Bcl-2 family members and the autophagy protein Atg6. In contrast, a variety of other oncogenic elements had no effect on MYC-VX-680 synthetic lethality. These findings indiate that MYC-VX-680 synthetic lethality is mechanistically distinct from MYC-primed apoptosis in response to a variety of apoptotic stimuli and that aside from elevated Bcl-2 family members, other pro-survival oncogenic lesions are unlikely to cause resistence to MYC-VX-680 synthetic lethality. In addtion, combined treatement with an AURKB inhibitor and small molecule inhibitors of the Bcl-2-family many prove a potent combination for treatment of MYC overexpessing tumors.

## Results

### A screening for modifers of MYC-VX-680 synthetic lethality

In RPE-MYC cells, VX-680 synthetic lethality is executed initially by a Bim-dependent apoptosis that kills approximately 30% of treated cells [5]. Cells that bypass apoptosis develop polyploidy, but these cells eventually succumb to a lethal form of autophagy that is dependent on Atg5, Atg6, and Atg7. This biphasic mode of cell killing by VX-680 enables study of the interplay between apoptosis and lethal autophagy and we set out to dissect molecular modifiers.

To model the gain or loss of cancer genes, we transfected and selected RPE-MYC cells using vectors expressing oncogenes and dominant negative forms of tumor suppressors. We exposed the resulting pools of puromycin resistant cells to 300 nM of VX-680 for three days, and quantified viable cells at 3 and 6 days after treatment. In this model system, the mode of death that occurs within the first 3 days (early death) is predominantly apoptotic. The mode of cell death that occurs at day 6 following release from VX-680 (delay death) is a lethal autophagic loss of the remaining polyploid population of cells (Fig. 1A).

**Figure 1.**
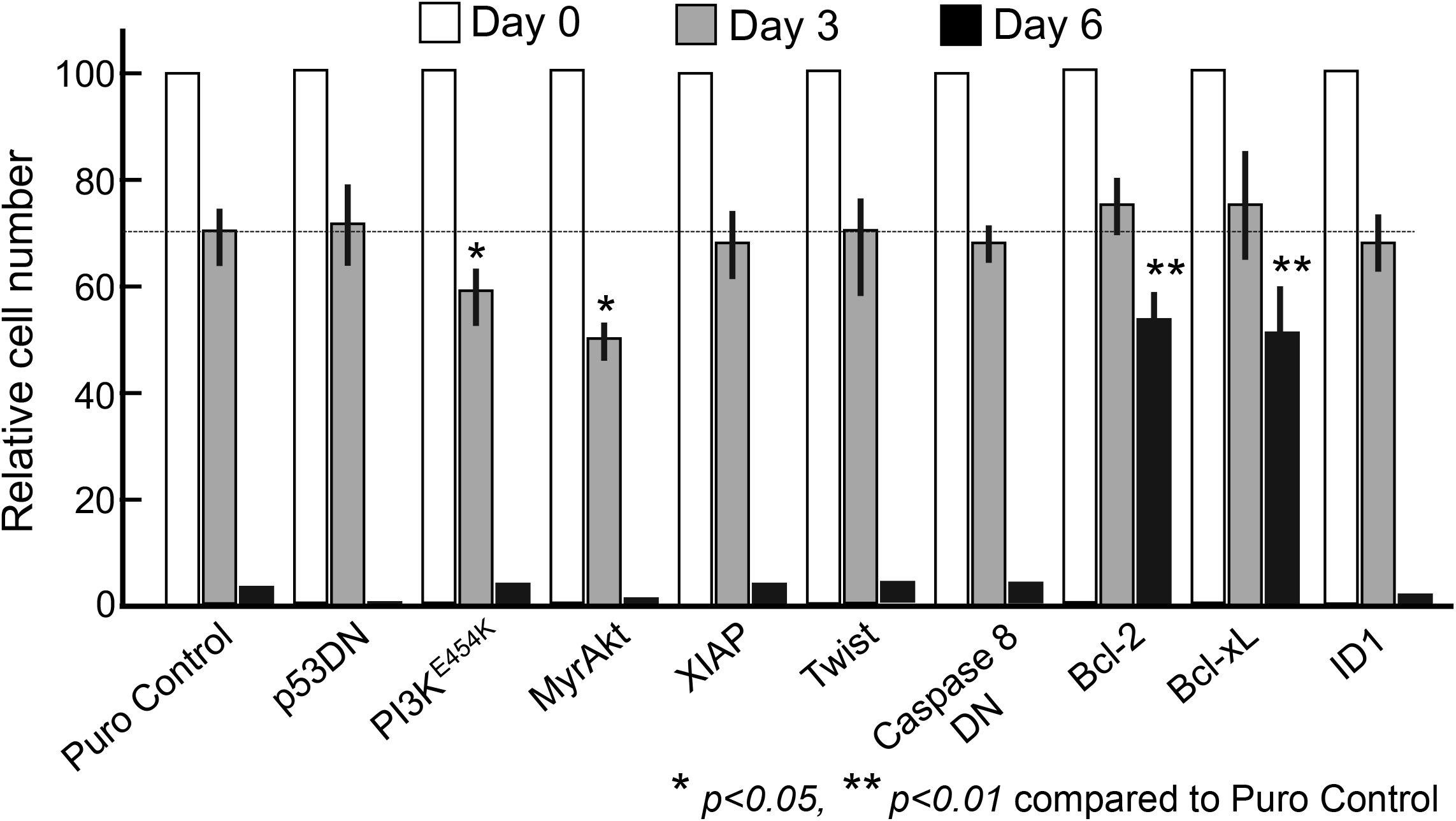
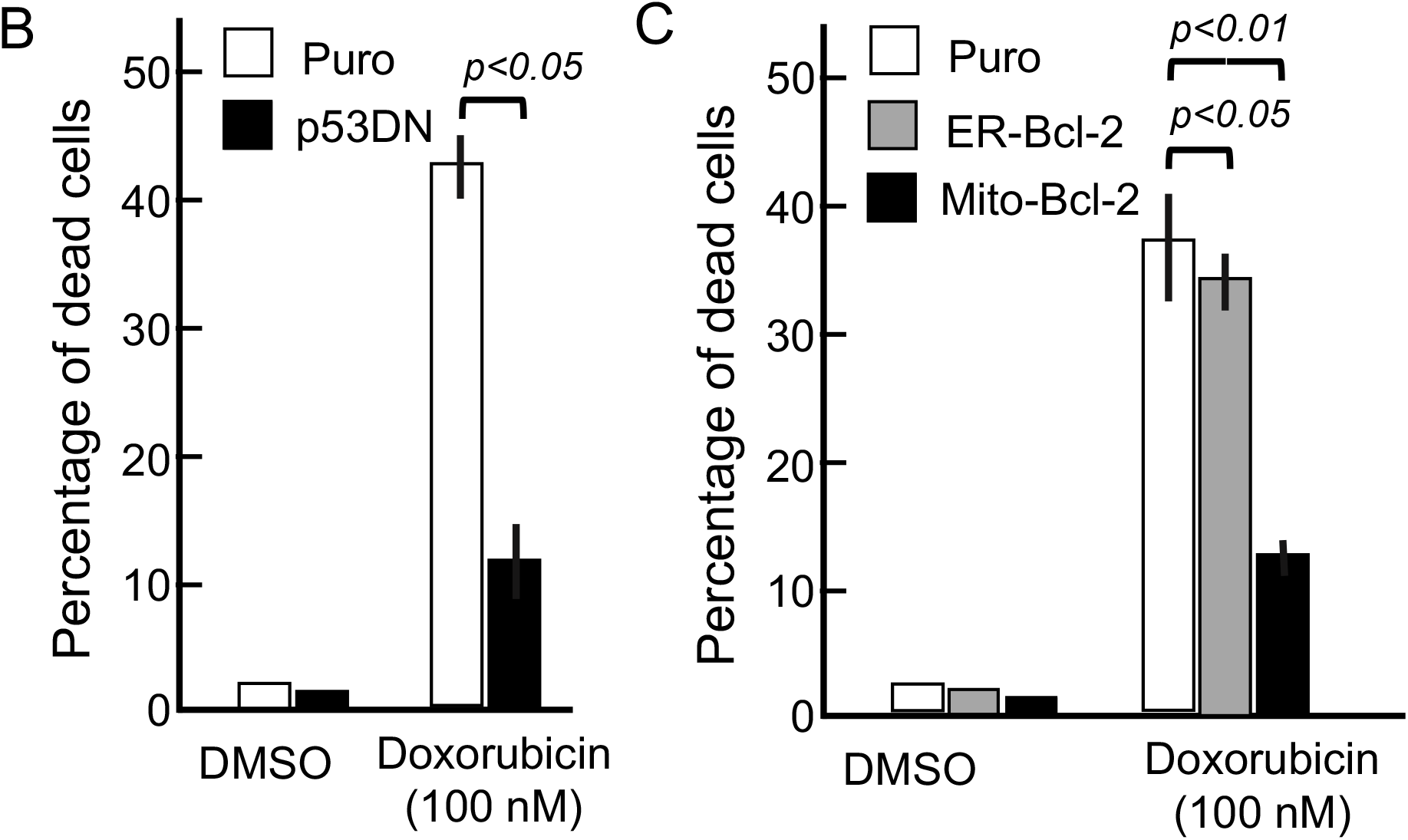
A screen for modifiers of MYC-VX-680 synthetic lethality identifies Bcl-2 and Bcl-xL as modifiers late, but not early, cell death. **A.** As a screen for modifiers of MYC-VX-680 synthetic lethality, RPE-MYC cells were stably transfected with either an empty vector control or a vector expressing the indicated genes and gene mutants. Cells were treated with 300 nM VX-680 for 3 days, and then transferred to drug-free medium. Cells were harvested at 0, 3 and 6 days after initiation of VX-680 treatment and assayed for viability by trypan blue exclusion assay. **B-C.** The indicated RPE-MYC derivatives were exposed to DMSO or 100 nM Doxorubicin for 2 days and then assayed for viability by trypan blue exclusion assay. **B.** p53DN diminishs apoptosis elicited by the DNA damaging agent doxorubicin. **C.** Mito-Bcl-2 not ER-Bcl-2 confers protection against apoptosis elicited by doxorubicin. Results in **A-C** represent the mean ± standard deviation for combined data from three independent experiments. Two tailed, unpaired Student’s T tests were used to derive *p* values.

In some contexts, MYC enhances a p53-dependent apoptosis in response to chemotherapeutics [12]. We interfered with p53 function using a dominant negative p53 mutant (p53DN) but this failed to protect RPE-MYC cells from VX-680-induced cell death (Fig. 1A). Expression of p53DN conferred more than 70% protection against the cytotoxicity elicited by doxorubicin (Fig. 1B), confirming that p53 function was compromised by this mutant. Therefore, p53 was not required for either early apoptosis or delayed lethal autophagy elicited by VX-680 in cells overexpressing MYC. VX-680 has been suggested to selectively kill p53 deficient cells [13], but this was not recapitulated in RPE cells with p53DN expression alone or in p53^−/−^ murine embryonic fibrobalsts [5]. Nonetheless, p53 loss-of-function does not alter the MYC-VX-680 synthetic lethality observed here.

Survival signaling through the RAS-PI3K-AKT signaling axis can synergize with MYC in eliciting cellular transformation and tumorigenesis [14]. We found that overexpression of an active H-RAS^G12V^ allele elicited profound senescence in RPE-MYC cells, preventing analysis in our sceen. However, we were able to generate RPE-MYC cells expressing myristoylated, constitutively active Akt (MyrAKT) or an activated PI3K^E454K^ allele. At day 3 of treatment, both conferred a moderate increase in RPE-MYC sensitivity to VX-680 compared to cells expressing MYC alone. All cells died at day 6 irrespective of the presence of MyrAKT or PI3K^E454K^ (Fig. 1A). Thus, active signaling from the PI3K-AKT pathway enhanced, rather than blocked, the early apoptosis elicited by VX-680 with no effect on the lethal autophagy of polyploid cells.

Cancer cells that overexpress MYC frequently inactivate Caspase 8 to escape apoptosis [15]. Expression of a dominant negative version of Caspase 8 (Caspase 8 DN) failed to rescue MYC-overexpressing cells from lethality elicited by VX-680 (Fig. 1A). This finding is consistent with a report that VX-680 can effectively elicit apoptosis in cancer cell lines harboring deletion of Caspase 8 [16]. Similarly, ectopic expression of TWIST, ID1 and even XIAP, which should sequester and subsequently inactivate active caspases [17], failed to have an effect on MYC-VX-680 synthetic lethality (Fig. 1A).

In contrast to the above findings, we found that overexpression of either Bcl-2 or Bcl-xL, provided near 50% protection against the cytotoxicity of VX-680 six days after administration (Fig. 1A). Neither provided appreciable protection within the first three days of treatment (Fig. 1A). This was not due to defective expression as doxirubicin-induced cell death, as opposed to VX-680, could be blocked by either mitochondrial or edoplasmic reticulum targeting of Bcl-2 (Fig. 1C). In addtion, by day 6, all live cells had become multinucleated and polyploid (Fig. 2). This means that Bcl-2 and Bcl-xL rescued delayed autophagic cell death, rather than MYC-VX-680 early apoptosis.

**Figure 2.**
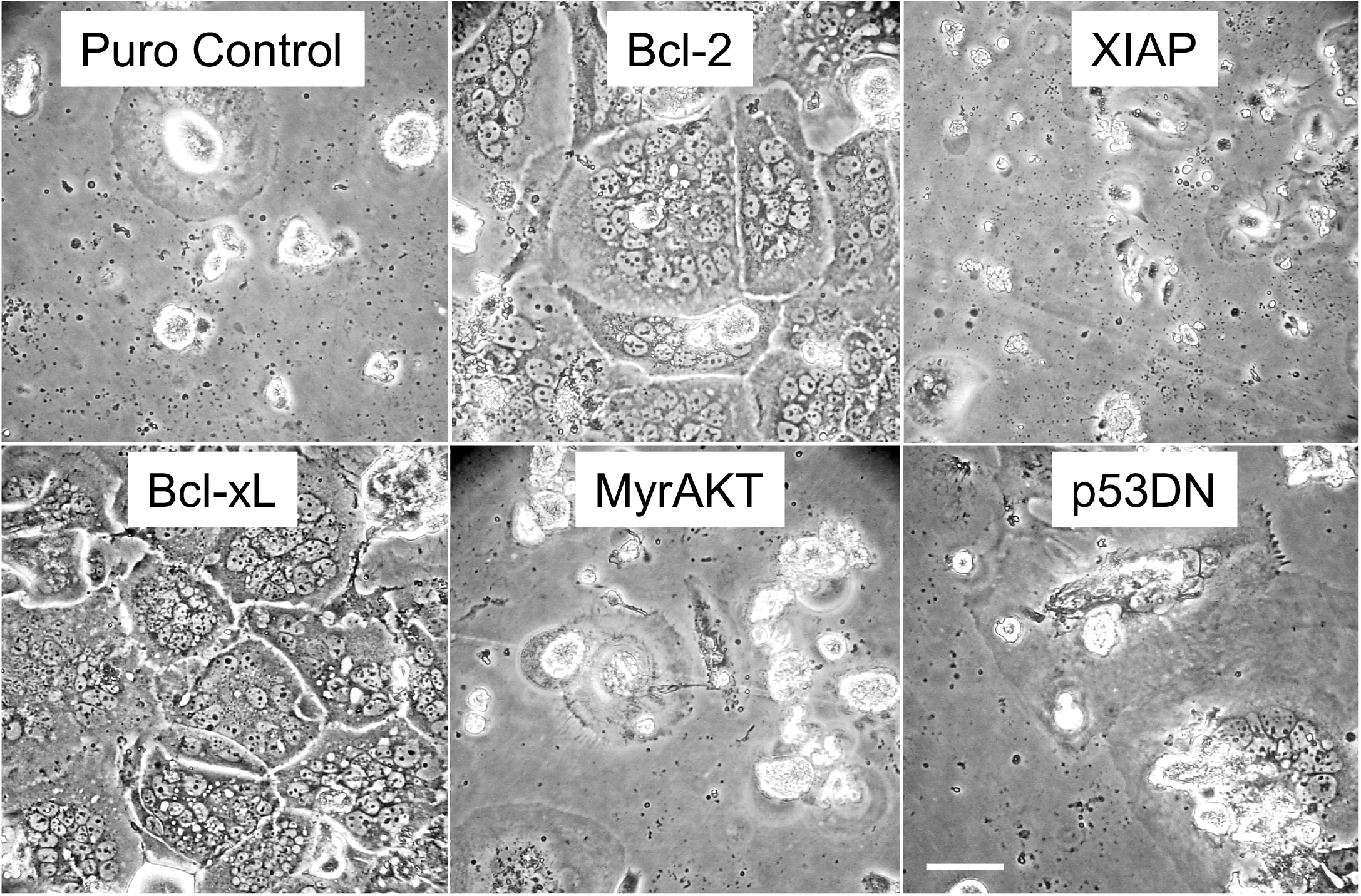
Bcl-2 and Bcl-xL promote the survival of polyploid cells. Light microscopy images of RPE-MYC cells that were stably transfected with either an empty puromycin control vector or a vector expressing the indicated genes. Cells were treated with 300 nM VX-680 for 3 days, and then transferred to drug-free medium. Micrographs were taken 6 days after initiation of VX-680 treatment. Scale bars are 50 μm.

Additional conclusions we can draw from our screen are first, that the intended target, AURKB, was successfully inhibited by VX-680 and that this inhibition was not influenced by any of our manipulations. This is because all of the treated cells developed polyploidy in response to VX-680. Second, the early phase of VX-680-induced apoptosis in RPE-MYC cells seems mechanistically distinct from previously reported MYC-dependent apoptosis in response to hypoxia, growth factor deprivation, and treatment with genotoxic chemotherapy. Finally, the genes and pathways without effect in our assay seem unlikely to cause resistence to MYC-VX-680 synthetic lethality, despite their frequent association with overexpression of MYC in human maligancies.

### Bcl-2 rescues from VX-680-induced death of polyploid RPE-MYC cells through interaction with Atg6

Becline 1/Atg6 is essential for the autophagic death of polyploid cells elicited by VX-680 [5] [18] and Bcl-2 and Bcl-xL have been implicated in regulation of autophagy through interaction with Beclin 1/Atg6 [19]. To further ellucidate how Bcl-2 and Bcl-xL protect polyploid cells, we tested a variety of mutants that are deficient in either inhibition of apoptosis or autophagy (Fig. 3A, B). Expression of the Bcl-xL mutant F131V, D133A, which is defective in binding to pro-apoptotic Bcl-2 family members Bax and Bak [20], failed to protect against early apoptotic cell death elicited by VX-680 (Fig. 3B, day 3). In contrast, this Bcl-xL mutant suppressed death of multinucleated cells (Fig. 3B, C, day 6 measurement), indicating that Bcl-xL interactions with pro-apoptotic Bcl-2 proteins are not required to rescue multinucleated cells from death.

**Figure 3.**
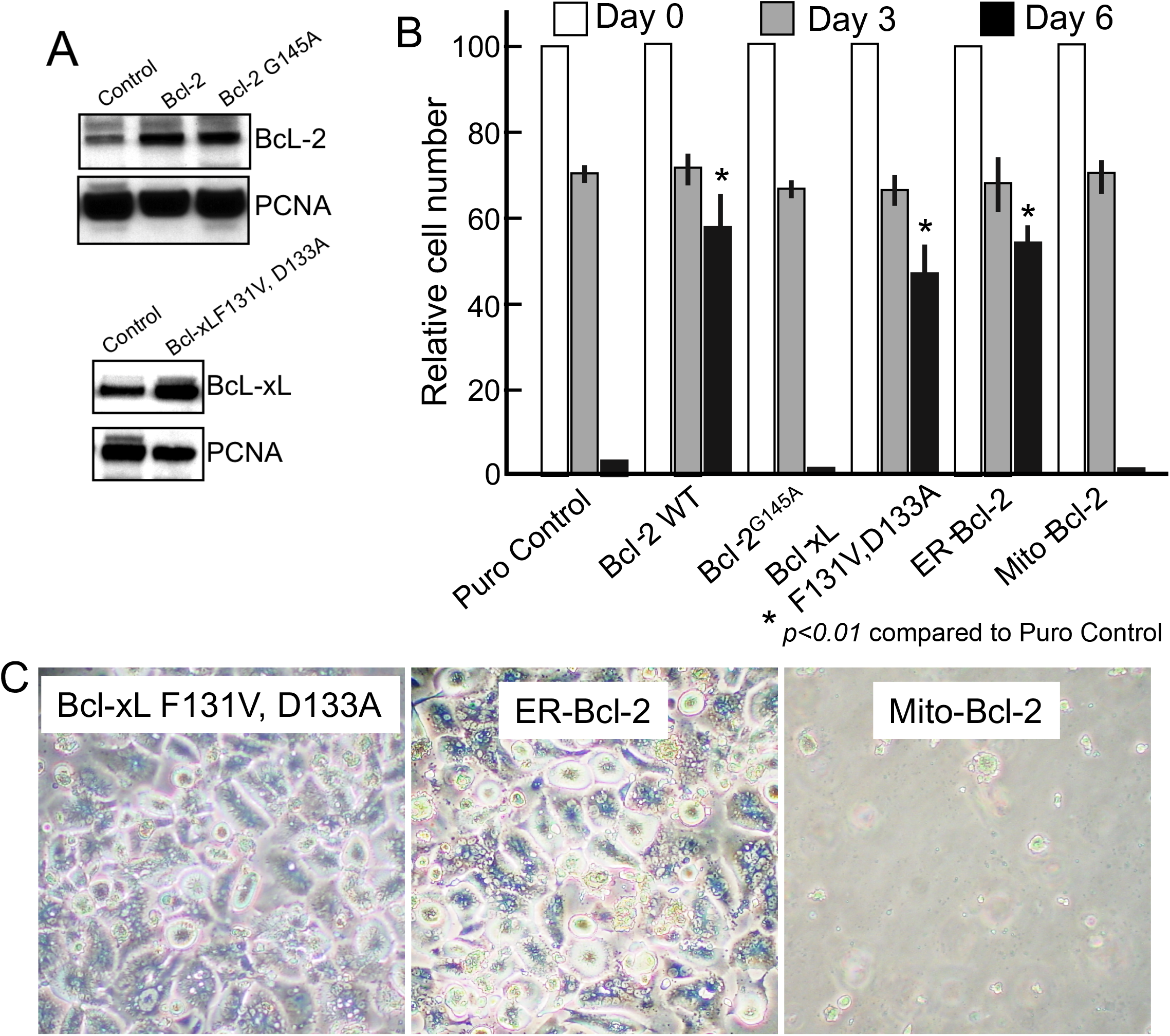
Bcl-2 and Bcl-xL promote the survival of polyploid cells through interactions that occur at the endoplasmic reticulum. **A.** Western analysis of whole cell lysates from RPE-MYC cells stably transfected with a control vector, wild-type Bcl-2, Bcl-2 G145A or Bcl-xL F131V, D133A. PCNA expression was used as a loading control. **B-C.** Differential inhibition of autophagic cell death by antiapoptotic members of the Bcl-2 family and their mutants. Cells were treated with 300 nM VX-680 for 3 days, and then transferred to fresh, drug-free media. Cells were harvested at 0, 3 and 6 days after initiation of VX-680 treatment and assayed for viability by the trypan blue exclusion assay. Results in **B** are expressed as a percent of the relative number of live cells at day 0 and represent the mean ± standard deviation for combined data from three independent experiments. Two tailed, unpaired Student’s T tests were used to derive *p* values. **C.** Light micrographs taken 6 days following initiation of VX-680 treatment (3 days after switch to fresh, drug-free media); magification 400X.

Next, we ectopically expressed a Bcl-2 mutant that fails to bind to Atg6, Bcl-2 G145A [19]. This mutant was unable to rescue from autophagic cell death (Fig. 3B), despite similar level of expression to its wild-type counterpart (Fig. 3A). Therefore, direct interaction with Atg6 was required for Bcl-2 to suppress MYC-VX-680 autophagic cell death. The Bcl-2 G145A mutant also had no apparent effect on early apoptosis elicited by VX-680 in RPE-MYC, despite previous reports that this mutant could function as a dominant negative form through dimerizing with endogenous Bcl-2 [21]. Nonetheless, we can concude from our assay that Bcl-2/Bcl-xL interactions with Atg6 are required to protect polyploid cells from MYC-VX-680 autophagic cell death.

### Endoplasmic reticular, rather than mitochondrial, Bcl-2 rescues VX-680-induced polyploid RPE-MYC cells

Bcl-2 is best known to exert anti-apoptotic function by interacting and neutralizing the pro-apoptotic members of the Bcl-2 family, such as Bax, Bak and Bim at the mitochondria [22]. However, suppression of autophagy by Bcl-2 requires its colocalization at the endoplasmic reticulum (ER), as opposed to mitochondria [19].

We tested whether either of these two subcellular locations for Bcl-2 were absolutely required to protect multinucleated cells from MYC-VX-680 synthetic lethality. We utilized two well characterized Bcl-2 fusion proteins, ER-Bcl-2 and Mito-Bcl-2. ER-Bcl-2 possesses an ER-targeting sequence from cytochrome b5 that enables ER-specific localization, while Mito-Bcl-2 harbors sequence from ActA of *Listeria monocytogenes* that functions as a guide to localize this fusion to the outer mitochondrial membrane [23]. Our findings with ER-Bcl-2 mirrored wild type Bcl-2 expression in preventing the delayed death but not early apoptosis in VX-680 treated RPE-MYC (Fig. 3B, C). In contrast, Mito-Bcl-2 was not able to rescue in this assay. We have shown that Mito-Bcl2 could protect against apoptosis elicited by the DNA damage agent Doxorubincin (Fig. 1C), confirming expression of functional mitochondrial protein. Therefore, we conclude that ER localized Bcl-2 is required to protect polyploid RPE-MYC cells after VX-680 treatment.

### Suppression of polyploid cell death by Bcl-2/Bcl-xL is associated with reduced autophagy

We carried out assays to definitively confirm inhibition of autophagy was the mode by which Bcl-2 and Bcl-xL protected polyploid cells from MYC-VX-680 synthetic lethality. During autophagosome maturation, soluble LC3-I (also referred to as MAP1LC3A or ATG8) is converted into its vesicle-bound form LC3-II by conjugation to phosphatidylethanolamine (PE) [24]. LC3-II puntate immunohistochemical staining marks autophagic cells. The same versions of Bcl-2 and Bcl-xL that were active in peventing death of multinucleated cells, also inhibited formation of LC3-II puntate staining. In contrast, mutants that did not prevent MYC-VX-680 synthetic lethality had ample puntate LC3-II staining, indicative of autophagy (Fig. 4A).

**Figure 4.**
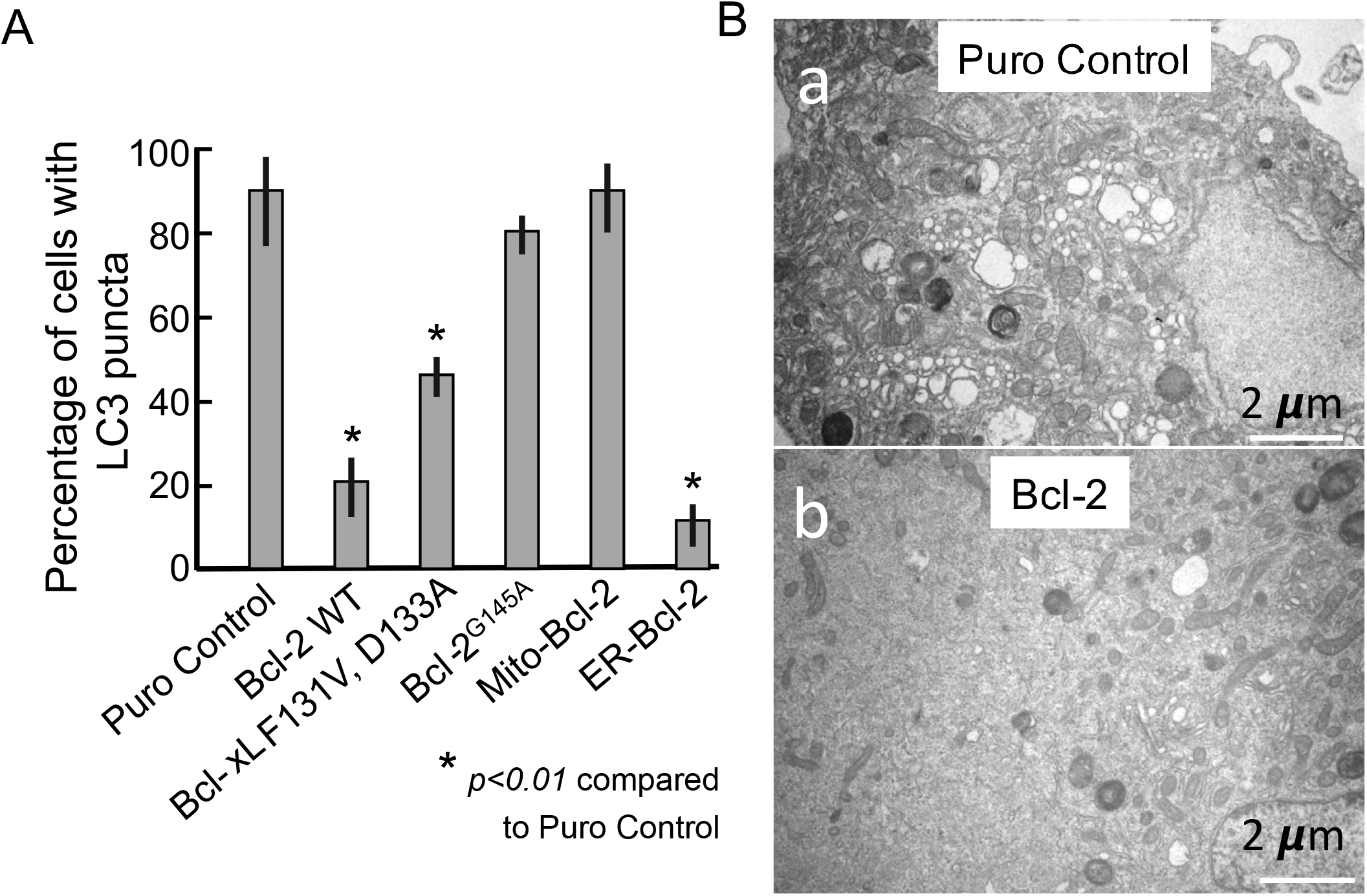
Bcl-2 and Bcl-xL inhibit autophagy in polyploid cells elicited by MYC-VX-680 synthetic lethality. **A.** Quantification of LC3-positive autophagic vesicles in RPE-MYC cells that were stably transfected with vectors expressing the indicated genes. The RPE-MYC derivatives were treated with 300 nM VX-680 for 3 days and then transferred into drug-free medium. As a surrogate for autophagy we quantified Atg8/LC3 antibody positive foci (puncta), 5 days after initiation of VX-680 treatment. Cells that displayed predominantly punctate Atg8 immunostaining and had more than 10 such puncta per cell were scored as positive. Two tailed, unpaired Student’s T tests were used to derive *p* values. **B.** Ultrastructural analysis of cell morphology. RPE-MYC derivatives were treated with 300 nM VX-680 for 3 days and then transferred into drug-free medium. Representative electron micrographs are shown for RPE-MYC-puro control cells (a) and RPE-MYC cells expressing Bcl-2 (b), 6 days after administration of VX-680 (Scale bar, 2μm).

Transmission electron microscopy (TEM) to examine cells for double membrane vesicles is the gold standard for detection of autophagy. TEM analysis confirmed that, when wild-type Bcl-2 was overexpressed, VX-680-induced autophagic vesicles were sparce (Fig. 4B, a vs. b). Collectively, these data indicate that polyploid RPE-MYC cell rescue by Bcl-2 and Bcl-xL requires an anti-autophagic activity, rather than anti-apoptotic activity. Rescue coincides with suppression of autophagy.

### Suppression of death of multinucleated cells by Bcl-2/Bcl-xL is associted with enhanced recovery after release from treatment

We sought to test whether protection of polyploid cells by Bcl-2 correlated with potential for therapeutic resistance. We perfomed long-term colony formation assays after drug treatment, which is one mode to assess the long-term effect of therapeutics on tumor cells. We found that rare polyploid RPE-MYC cells can resume proliferation and reverted back to mononucleated cells with cellular and nuclear size and morphology similar to that of parental cells. Out of one million polyploid cells, about 10 cells resumed proliferation and formed a vialble colony (Fig. 5). We found that despite rescue of polyploid cell death at day 6 after VX-680 treatment, overexpression of Bcl-2 enhanced the long-term colony formation only 3-fold. Therefore, most polyploid cells still failed to renew proliferation. This indicates that VX-680, although largely effective against cells that overexpress MYC, may rarely be bypassed by manipulations that disrupt the induction of autophagy. It remains to be seen if this mechanism could contribute to resistance to synthetic lethal therapy beteween MYC and disabling the chromosomal passenger protein complex *in vivo*, but the data presented here support the evaluation of Bcl-2 family inhibitors alongside Aurora kinase inhibition.

**Figure 5.**
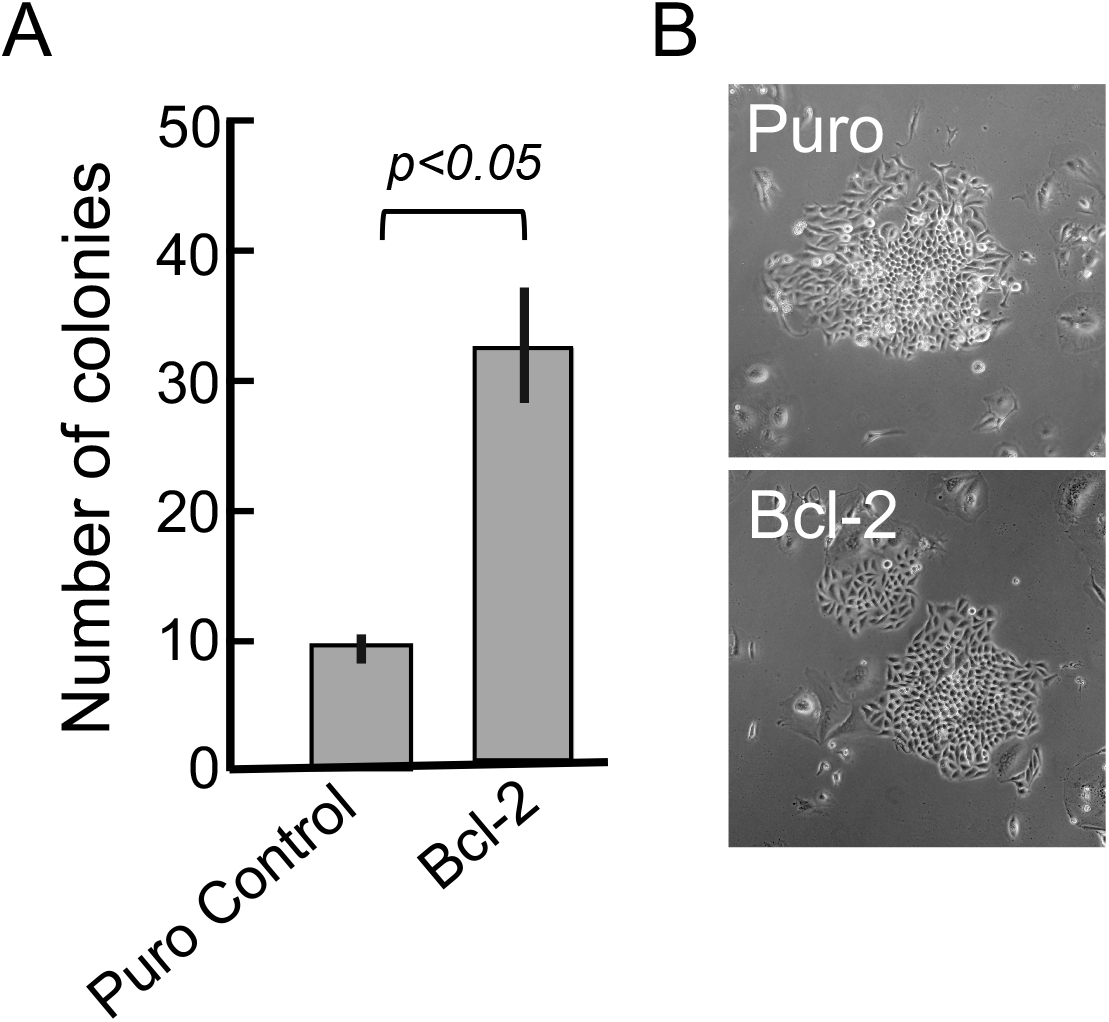
Bcl-2 and Bcl-xL enhance the long-term recovery of polyploid cells. RPE-MYC-puro cells or RPE-MYC expressing ectopic Bcl-2 were exposed to 300 nM of VX-680 for 3 days and then transferred to drug-free fresh medium. **A.** At day 8 after the release from the treatment with VX-680, colonies were ennumerated. **B.** Light micrographs of colonies from each group. Final colony and cell morphology was similar for each RPE-MYC derivative. Data in **A** represents the mean ± standard deviation for combined data from three independent experiments. Two tailed, unpaired Student’s T tests were used to derive *p* values.

## Discussion

Genetic events that occur in cancer can impart unique vulnerabilities on cancer cells that are not present in normal somatic cells. The promise of synthetic lethal approaches is to identify these unique vulnerabilities and drug these dependencies to the detriment of the cancer cell, while sparing normal cells. This is an especially attractive tactic with respect to oncogenic lesions that are difficult to target with current approaches.

MYC overexpressing tumors can be targeted through a synthetic lethal interaction with VX-680, a pan-Aurora kinase inhibitor [5]. Here, we undertook a systematic survey to screen for tumor associated genetic chages and pathways that could modify MYC-VX-680 synthetic lethality. We modeled each of the oncogenic alterations individually in RPE-MYC cells and tested this panel of isogenetic cell lines for their sensitivity to VX-680. One benefit to this experimental approach is the abiltiy to examine two distinct types of VX-680-induced death, early apoptotic death and later death of polyploid cells. This later death being associated with an apparently detrimental autophagy.

In contrast to the well established effect that many genetic events have on the inhibition of MYC-primed apoptosis [5] [6] [7], none of the tested manipulations in our assay enabled resistance to MYC-VX-680 early apoptosis. Our findings in these isogenic cells is congruent with previous studies in cancer cell lines suggesting MYC-VX-680 synthetic lethality still operates in cells with inactivation of p53 or Caspase 8 [15] [25]. The robustness and inability to challenge this apoptosis with traditional inhibitors of cell death suggests that the early apoptotic phase of MYC-VX-680 synthetic lethality may be mechanistically distinct from MYC-induced apoptosis in other systems. It suggests that mechanisms leading to resistance to tranditional DNA-damaging chemotherapy may not functionally oppose the early stages of MYC-VX-680 synthetic lethality.

In contrast to the lack of modifiers for early apoptosis in our screen, we found that Bcl-2 and Bcl-xL can protect from the late stages of MYC-VX-680 synthetic lethality. At this stage, polyploid cells predominate and these cells engage a cell death characterized by excessive autophagy. By day 9 of culture, even polyploid cells are cleared from culture. The pattern of early and late cell death with VX-680 is observed in other MYC expressing cells, including mouse embryo fibroblasts (MEF-MYC), rat embryo fibroblasts (Rat1A-MYC) and 14 different human cancer cell lines, so it is not unique to MYC expressing RPE cells [5].

Our observations with Bcl-2 and Bcl-xL are consistent with findings that Bcl-2 inhibitors and the AURKB inhibitor AZD1152 act synergistically to kill tumor cells [26] [27]. However, this effect has been attributed to apoptotic elimination of polyploid cells in the presence of Bcl-2 inhibitors. We have uncovered a distinct mechanism. In our assays the anti-apoptotic members of the Bcl-2 family work independent of mitochondrial apoptosis. Our mutant studies suggest Bcl-2 and Bcl-xL work though interaction with Beclin-1/Atg6 at the endoplasmic reticulum to reduce autophagy and the second phase of MYC-VX-680 synthetic lethal cell death.

Our findings reveal that Bcl-2 and Bcl-xL are determinants of polyploid cell viability following aurora kinase inhibition in MYC overxpressing cells. It is unknown if the increase we observed in long-term colony formation could correlate with drug resistance in humans, but these observations do suggest a mechanism by which resistance to MYC-VX-680 synthetic lethality might arise through interferance with induction of lethal autophagy. Therapeutic escape of even a few cancer cells could have an enormous effect on patient outcome. BH3-mimetic drugs, such as Venetoclax and Navitoclax, are able to inhibit the interaction between Bcl-2 and Atg6 [28]. Our findings lend weight to the argument that these Bcl-2 inhibitors combined with chromosomal passenger protein complex inhibition could be a potent combination therapy for tumors that overexpress MYC.

## Materials and Methods

### Chemicals, cell lines and culture media

VX-680 was purchased from Selleck Chemicals, validated by analytic HPLC in-house, and used at concentrations indicated in figures and figure legends. Generation of RPE-NEO and RPE-MYC were described before [6]. Sources for some of the mutant genes utilzed are as follows: vector expressing dominant negative p53 (Takara); myristoylated AKT (pMIG-myrAKT, Addgene); mutant PI3K p110α subunit (pMIG-PI3KE545K, Addgene); Bcl-2 proteins and their mutants (pMIG-Bcl-xL from Addgene, pMIG-BCL2 [6], pcDNA3.1-BCL2 G145A [19], pMIG-Bcl-xL F131V, D133A [20], Bcl-2-cb5 [23] Bcl-2-acta [23]). The identitiy of plasmids was confirmed by DNA sequencing. For generation of expression pools, RPE-MYC cells were cotransfected with expression vectors and pMSCV-puro followed by puromycin selection (1 μg/ml) to generate their derivatives. RPE-MYC Puro control cells were generated by transfection with pMSCV-puro only. All cell lines were cultured in DMEM supplemented with 5% fetal bovine serum and at 5% CO_2_ and 95% O_2_ in a humidified incubator.

### Cell viability and colony formation assays

Cells were seeded to 24-well plates and allowed to adhere overnight at 37°C before drug treatment. For quantitation of viability, cells were washed with PBS once, treated with 0.25% trypsin-EDTA solution, resuspended in DMEM and counted with a hematocytometer under either a light microscope or Luna™ cell counter after staining with trypan blue (0.04%, 1:1). Colony formation assays were carried out as described [5]. Briefly, cells at 20% confluence in 6-well plates were exposed for 3 days to 300 nM of VX-680, transferred to drug-free fresh medium and then cultured for another 7-10 days before staining with crystal violet to visualize colonies.

### Western analysis

Whole cell extracts were prepared by incubating cells for 15 minutes at 4°C in lysis buffer [50 mM Tris (pH 7.5), 200 mM NaCL, 0.1% SDS, 1% Triton X-100, 0.1 mM DTT, and 0.5 mM EGTA] supplemented with protease inhibitor mixture (BD Biosciences). The extracts were centrifuged at 8,000 × g for 10 min to clear insoluble material. Protein concentration in the supernatant was determined using a Bio-Rad protein assay. Lysate (50–100 μg protein) was resolved on NuPAGE (4–12%) Bis-Tris gels (Invitrogen) and transferred to nitrocellulose membranes (Bio-Rad). Blocking was with 5% nonfat milk in PBS buffer for 1 hour. Membranes were incubated overnight at 4°C with primary antibodies diluted 1:1,000 in blocking buffer. Rabbit polyclonal antibodies were used for Bcl-2, Bcl-xL and PCNA. Horseradish peroxidase-conjugated anti-rabbit immunoglobulins were from Santa Cruz Biotechnology. Western blots were developed with the SuperSignal West Femto or Pico ECL detection kit (Thermo Scientific).

### Fluorescence and transmission electron microscopy

Immunofluorescence staining using a mouse monoclonal antibody against LC3 has been described [5]. Cells were cultured on coverslips in a 6-well plate, fixed with 4% paraformaldehyde and then permeabilized with 0.3% Triton X-100. A mouse monoclonal antibody (clone 5F10) from NanoTools was used to detect endogenous Atg8b/LC3. Primary antibodies were detected with Texas red-conjugated or fluorescein isothiocyanate-conjugated secondary antibodies (Jackson ImmunoResearch). After immunostaining, we mounted cells on microscope slides with 4’,6’-diamidino-2-phenylindole (DAPI)-containing Vectashield mounting solution (Vector Laboratories). For fluorescence detection, we used an EVOS FL fluorescence microscope (ThermoFisher). For transmission electron microscopy, cells were fixed at 4 °C in 0.1M Cacodylate buffer (pH.7.4) containing 3% glutaraldehyde and 1% paraformaldehyde and then embedded in Eponate 12. Electron microscopy was performed by the Microscopy and Advanced Imaging Core Facility, San Francisco VA Medical Center, with a Tecnei 10 transmission electron microscope at 80 kV on ultrathin sections (80 nm).

### Statistical analysis

Statistical analyses were performed with the GraphPad Prism software. Statistical significance was evaluated with Student’s unpaired, two-tailed *t* test. *P* values less than 0.05 were considered statistically significant.

## Acknowledgements

We thank Beth Levine for the pcDNA3.1-Bcl2-G145A construct, Stanley Korsmeyer for pMIG-Bcl-x_L_ F131V, D133A, David Andrews for Bcl-2-cb5 and Bcl-2-acta. We thank Ivy Hsieh and Juan Engel at San Francisco Veterian Affairs for excellent technical support with transmission electron microscopy. This work was initiated at J. Michael Bishop Laboratory at University of California, San Francisco with a funding support from the G. W. Hooper Research Foundation and was completed at J. Michael Bishop Institute of Cancer Research in Chengdu, China, with an institute endowment fund.

